# Task-Free Recovery and Spatial Characterization of a P3-Like Global Network from Resting-State EEG

**DOI:** 10.1101/2020.09.27.316141

**Authors:** Akaysha C. Tang, Adam John Privitera

## Abstract

Diagnosis of mental illness, testing of treatment effects, and design of prevention strategies all require brain-based biomarkers that can serve as effective targets of evaluation. The search for such markers often starts with a search for neural correlates from brain imaging studies with measures of functions and behavior of interest. Yet such an approach can produce erroneous results for correlations do not guarantee causation. Only when the markers map onto neurocomputationally-relevant parameters can such markers best serve the intended function. Here we take an alternative approach to begin with targeting the neuroanatomically and neurophysiologically well-defined neuromoduatory systems that are well positioned to serve the computational role of generating globally synchronized neural activity for the purpose of functional integration [1]. By applying second-order blind identification (SOBI) [2], a blind source separation algorithm (BSS), to five minutes of resting-state EEG data (n=13), we provide evidence to support our conclusion that neuroelectrical signals associated with synchronized global network activity can be extracted using the detailed temporal information in the on-going continuously recorded EEG, instead of event-related potentials (ERPs). We report reliable extraction of a SOBI component, which we refer to as the P3-like component, in every individual studied, replicating our earlier report on data from a single participant [3]. We show that individual differences in the neural networks underlying this P3-like component can be revealed in high dimensional space by a vector of hits-based measures [4] for each of the P3-like network’s constituent structures. Given that resting-state EEG can be obtained with greater ease at natural non-hospital settings and at much lower cost in comparison with fMRI, and that mobile EEG systems have become increasingly available, the present work offers an enabling technology to support rapid and low-cost assessment of much larger and diverse populations of individuals, addressing several methodological limitations in our current investigation of brain function. Future opportunities and current limitations will be discussed.

## I. Introduction

Three major reasons have been identified for why neurobiology has yet to effectively impact mental health practice [5, 6] which we understand as the following. The first is the nearly exclusive reliance of these studies on the research framework of group comparisons, as in the case of comparing diseased populations with healthy controls or two different healthy populations. The second is the associated lack of sufficient attention to how to provide critical and meaningful information about the status and outcomes of a given individual. The third is the lack of identification of neuromarkers based on neurocomputationally relevant parameters. Here, building upon decades of effort on EEG-based source imaging [7-13], we present an EEG-based source-imaging study of the resting-state brain to show that a neurobiologically-grounded and network configuration-based characterization can be provided for each individual without using group data, and that cross-individual variability as well as consistency in the involvement of each brain region in a globally synchronized cortical network can be quantitatively characterized.

In search of neurocomputationally relevant parameters, we capitalize on the extensive psychophysiological and neurobiological literature on the neuromodulatory systems underlying novelty and uncertainty detection and their well-documented expression in the form of the EEG-derived P3 component [1, 14-17]. Ample neurobiological evidence indicates that the spatially most extensive and globally synchronized innervation of the entire cortical mantel is provided by neuromodulatory systems. Cholinergic and noradrenergic mediated global modulation are among the most extensively studied. These systems consist of projection neurons from subcortical structures to the cerebral cortex, and are thus capable of producing temporally synchronized changes in neuronal excitability and synaptic transmission across the entire cerebral cortex.

Recent applications of second-order blind identification (SOBI), a blind source separation algorithm, to EEG data offer evidence that such global synchronizations can be extracted from EEG. SOBI was able to recover, from different tasks conditions (oddball as well face perception tasks), the P3 component whose scalp projection and subsequent source location all revealed a globally distributed network involving the frontal, temporal, parietal, and occipital lobes [4, 18, 19]. The recovery of these SOBI P3 components are as expected in principle because SOBI works by using temporal information in the ongoing EEGs, not averaged ERPs computed using task parameters. Essentially, signals originating from different brain structures but with identical time courses will be separated from other functionally distinct signals into a single component. Thus, by definition, the activity of the SOBI-recovered P3 component reflects synchronized neuroelectrical activity across all constituent parts of the whole underlying network.

Because SOBI works by using cross-correlations from the continuous EEG data instead of using event related information, it should be able to recover source signals associated with functionally distinct neural networks without requiring the use of task related information, such as event-related potentials (ERPs). Indeed, our earlier case study [3] partially confirmed this prediction. Among the components recovered by SOBI from a few minutes of resting-state EEG data (collected from a single individual attending a US university), we found one whose scalp projection pattern closely resembled that of the SOBI-recovered P3 component from EEG collected during a visual oddball task. This observation motivated us to consider this P3-like component as a potential novel neuromarker to capture the synchronized global activity.

The present study has the following aims: (1) replicating the earlier finding of extracting this P3-like component from resting-state EEG data by investigating a larger sample from 13 individuals attending a university in Hong Kong; (2) determining whether the SOBI-recovered P3-like components can be localized using similar methods used for localizing the SOBI-recovered P3 component; (3) providing quantitative characterization of the spatial configuration of the P3-like components using a recently introduced hits-based analysis method [4, 19].

## II. Methods

### A. Experimental procedures

Approval for this study was granted by the Human Research Ethics Committee of the University of Hong Kong. We report data from five minutes of resting-state EEG data collected from thirteen right-handed participants sitting quietly with their eyes closed (6 males) between 19-33 years of age (M = 26.50 ± 4.48 years) with no reported neurological conditions. Continuous reference-free EEG data were collected in an unshielded room using an active 128-channel Ag/AgCl electrode cap, ActiCHamp amplifier, and PyCorder data acquisition software (Brain Vision, LLC) with a sampling rate of 1000 Hz. Data collection began only after the impedance of all electrodes was below 10 KΩ. Before additional processing, a 50 Hz notch filter was applied to raw EEG data in order to remove line noise. Data were spatially down-sampled to 64-channel to allow for comparison with our previous work on localization [4, 18].

### B. Extraction of the P3-like component using second-order bind source separation (SOBI)

SOBI [2] was applied to continuous EEG data to decompose the *n*-channel EEG data into *n*-components (**Fig. 1, Step 1**), each of which corresponds to a recovered putative source that contributes to the scalp recorded EEG signals. Detailed descriptions of SOBI’s usage [3, 20-24], SOBI validation [25, 26], and review of SOBI usage [27-29] can be found elsewhere. For further details about the use of SOBI on continuous EEG data, please see the *Methods* section of our companion paper [4].

**Fig. 1.**
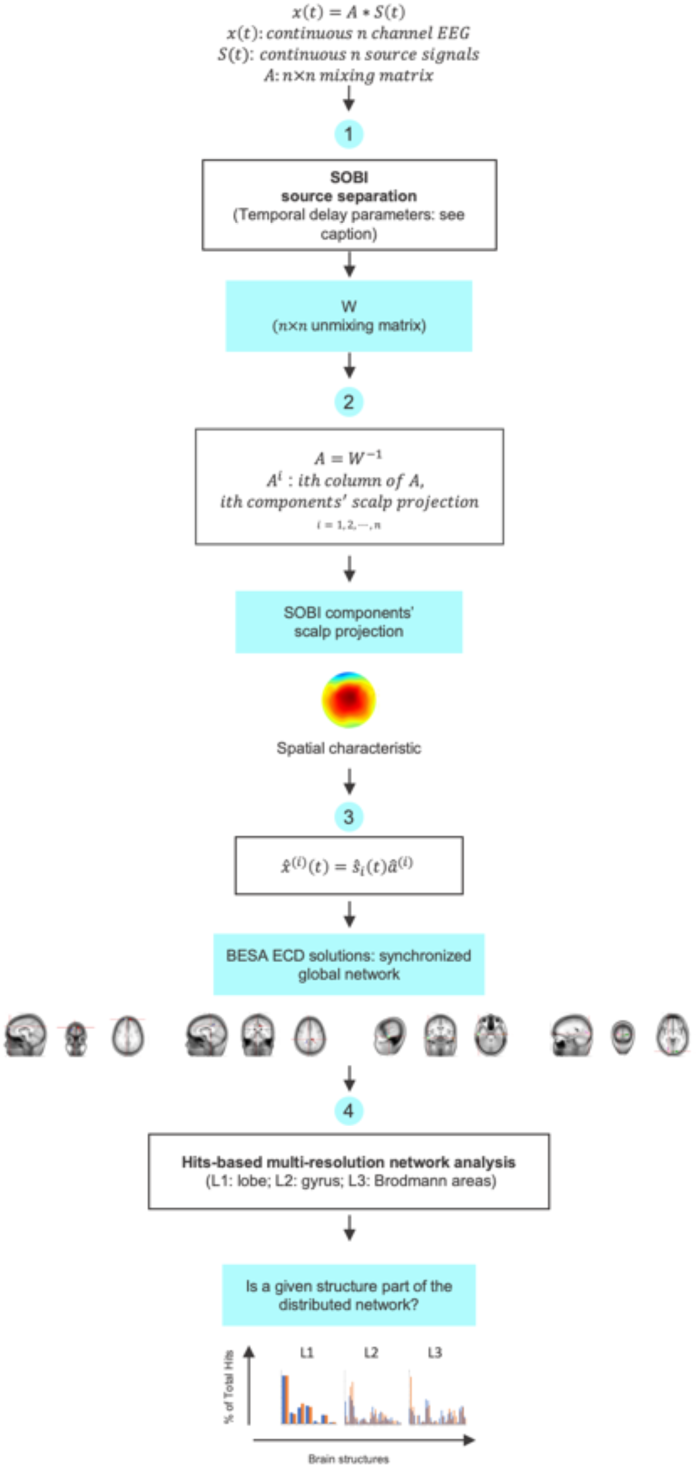
The analysis pipeline used for processing of individual data. **(1)** appication of SOBI to continuous EEG data, **(2)** identification of P3-like component using P3 component’s spatial characteristics, **(3)** localizing the generators of the P3-like component via equivalent current dipole (ECD) modeling, and **(4)** hits-based analysis of P3-like network configuration.

Here, SOBI-recovered P3-like components (hereafter, P3-like components) were identified according to spatial information alone given in the scalp projection pattern of the SOBI component derived from unmixing matrix W (**Fig. 1 Step 2)**. Here, the P3-like component is operationally defined by its scalp projection pattern displaying a bilaterally symmetric concentric dipolar field pattern similar to the scalp projection of the P3 component [4, 18].

Similar to the modeling of the SOBI recovered P3 components, the generators of this P3*-*like component were also modeled as a set of ECDs using BESA (BESA Research 6.1, Brain Electrical Source Analysis). From previous localization of SOBI-recovered P3 components, we know that such a global network typically consists of 4±1 pairs of symmetrically placed dipoles. Using this prior knowledge, we started the ECD modeling with a starting configuration, consisting of the same previously used ECD starting template, that is a model consisting of four pairs of bilaterally symmetrically placed dipoles at broadly-distributed locations roughly covering all four lobes of the cerebral cortex **(Fig. 2)**. For other details, see our companion paper [4].

**Fig. 2.**
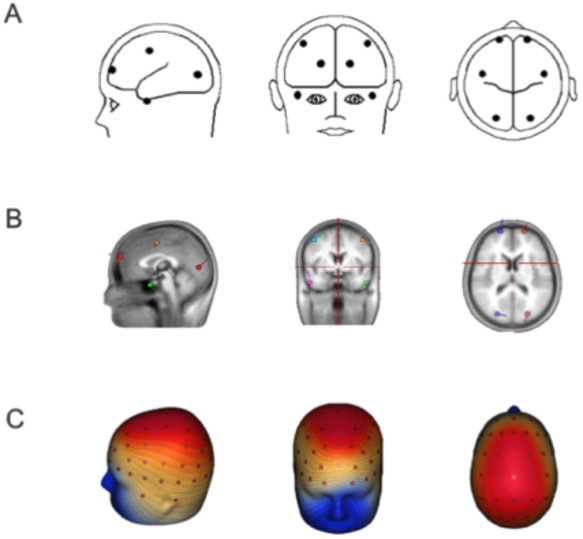
The starting dipole configurations used for the localization of the P3-like components in BESA. **AB**. starting tempate shown in schematic and sMRI views. **C**. scalp projection of the template.

Unlike data collected during an oddball task, resting-state EEG data do not afford any event-related potentials (ERPs), which are typically projected onto the scalp when localizating P3 components. In determining what could serve as alternative signals used as input to BESA, we note the fact that SOBI components’ scalp projections are time-invariant (see equation in **Fig. 1, Step 2)**. Therefore, the waveforms should not, in priciple, affect the localization solution. Therefore, to avoid numerical issues, a single ERP waveform from the P3 component of the same participant was used in the computation of **Fig. 1 Step 3**. The solution of the source location was given in dipole coordinates [30] and used as input to the following fits-based analysis.

### C. Analysis of hits associated with anatomical structures

Talairach Client (version 2.4.3) [31] takes BESA output as inputs and outputs structures and associated hits numbers at lobe, gyrus, and cell type levels. Similar to the analysis of the P3 components from the same set of individuals, we report the probability of each structure being observed as part of the P3-like network and the % of total hits that a given structure makes up in each individual’s total hits for the P3-like components. Statistical analyses were performed similarly to those for the P3 components.

## III. Results

### A. Reliable recovery of P3-like components from resting-state EEG using SOBI

Through visual inspection of all of the scalp projections of SOBI components recovered from individual resting-state EEG data, we were able to identify, in each of the 13 participants, at least one component whose scalp projection patterns mirrored those of the classic P3 components (the exact binomial test, *p* < 0.001). Similar to the P3 scalp topographies (**Fig. 3B**), these resting-state EEG derived P3-like scalp projections (**Fig. 3A)** all have bilaterally symmetric and nearly concentric scalp projection patterns, with variations in their centers along the midline, some centered over the central sulcus, and others more anterior or posterior. Note that although the precise topography of these components differs across individuals, the average scalp projections of the two components are rather similar. These results demonstrate that SOBI performs robustly in extracting the P3-like component from each individual’s EEG data.

**Fig. 3.**
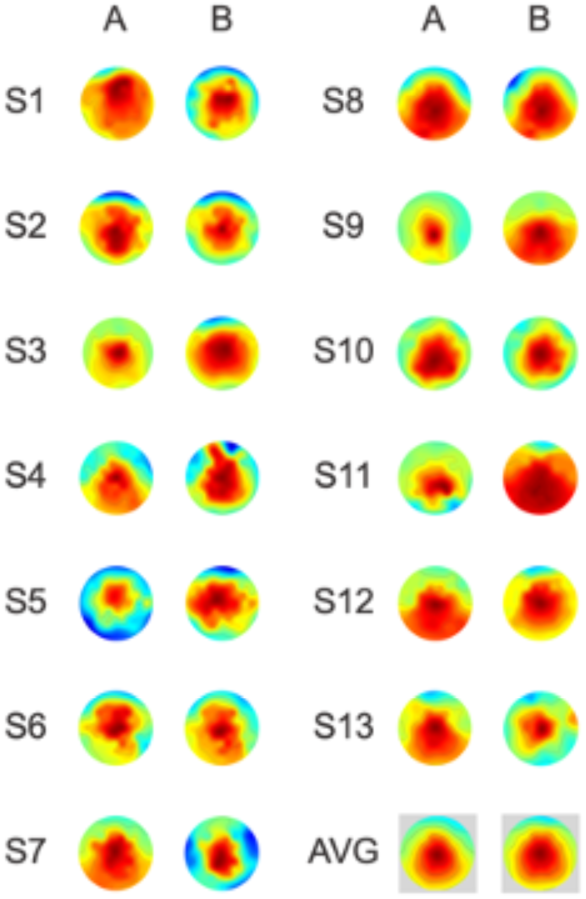
Reliable identification of P3-like components by SOBI and individual differences. Compare the scalp projections of the P3-like components (**A**: from resting EEG) with the P3 components (**B**: from EEG during a visual oddball task). Note the similarity between the average scalp projections despite cross-individual differences.

### B. Variable neural generators underlying the SOBI P3-like component

To determine the underlying neural networks that generate the P3-like components, the scalp topography of each individual’s component was modeled as a network of ECDs with an average goodness of fit (GoF) of 97.36 ± 0.4% (n=13). **Fig. 4A** shows an example from one individual, for whom, the scalp projection of the P3-like component (**Fig. 4A: Data**) is well fitted by the similar model scalp projection (**Fig. 4A: Model**) with little remaining unexplained variance (**Fig. 4A: Residual)**. The ECDs of the model are shown against the structural MRI of an average brain provided in BESA (**Fig. 4B)**.

**Fig. 4.**
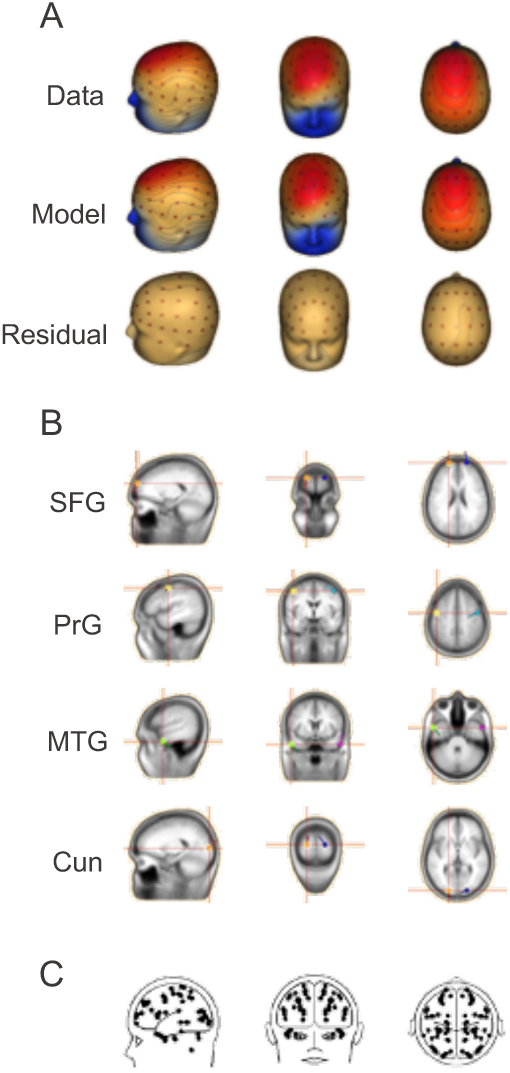
ECD modeling of SOBI-recovered P3-like components. **AB:** example from a single participant. **C**. overlaid ECD solutions from all individuals. **A**. Data: an individual’s P3-like component scalp projection; model: the ECD model projection; residual: the difference between data and model. **B**. ECD solution shown against the structural MRI (sMRI) of an average brain provided by BESA. **C**. Overlaid dipoles from all participants’ P3-like components showed no focal clustering.

Consistent with the variable 2D scalp projection patterns in **Fig. 3A**, the spatial distribution of the ECDs are also variable cross-individual, as indicated by the overlaid ECD solutions from all participants (**Fig. 4C)**. Because highly consistent patterns across individuals would result in clustered distribution of ECDs, the lack of clustering indicates a high degree of variation in the spatial configuration of the P3-like network. A similar broadly-distributed set of underlying neural generators were observed for a SOBI-recovered P3 component from visual oddball data [4], suggesting that both of these networks are similar in their variable configurations.

### C. Quantifying cross-individual variability in network configuration

The probability of a given brain structure’s involvement in the P3-like network was used to quantify the cross-individual variability. Hits were found in the frontal lobe in 100% of the participants studied (13 out of 13, exact binomial test, *p* < 0.001), in the parietal, temporal, and occipital lobes in 77% of the participants (10 out of 13, exact binomial test, *p* < 0.05), in the limbic lobe in 31% of participants (not significant), and in the cerebellum in 15% participants (not significant). The “gyrus” and “cell type” levels of analysis also show a greater cross-individual reliability for frontal structures than the remaining structures. At the “gyrus” level, although hits were found in 30 structures out of a possible 55, only MFG and SFG were observed reliably with statistical significance (77%, in 10 of 13 participants, *p* < 0.05). At the “cell type” level, while hits were found in 30 out of a possible 71 structures, none of the remaining structures are reliably involved across individuals, with BA9 and BA18 having the highest probability of being observed in 62% (8 of 13) of the participants.

These results show that the P3-like components consists of a combination of more reliable frontal lobe structures and other less consistently found non-frontal structures. The more consistent cross-individual involvement of the frontal lobe suggests that even under non-task conditions, the frontal lobe can still play a dominant role in this global network while the more variable involvement of other brain structures in this network may reflect individual differences in how the frontal lobe engages with the rest of the brain.

### D. Quantifying cross-individual variations in network configuration

That percentage of total hits that each of the constituent structure makes up was used to quantify the relative contribution of each structure towards the overall P3-like network. **Table 1** reveals at all three levels of analysis, the average % of total hits over all individuals and their *Z* statistics. Analysis of the % of total hits variable showed that 4 structures at the lobe level, 14 at the gyrus level, and 15 at the cell type level made statistically significant contributions to the whole P3-like network (Wilcoxon signed rank tests, 1-tailed). The highest % of total hits measures were 45% for the frontal lobe: 13% for MFG, and 11% for both BA6 and BA8.

**Table 1.** Hits analysis of P3-like network at the Lobe **(A)**, Gyrus **(B)**, and Cell Type **(C)** levels reveal large overlap between the P3-like and P3 networks. Nhits: the number of hits associated with a structure. % of total hits: average % of total hits per structure (n=13). *Z* and *p*: statistics from one-sample Wilcoxon signed-rank test (1-tailed). Shared structures between the P3-like and P3 networks are highlighted.

| A Lobe Level |  |  |  |  |  |
| --- | --- | --- | --- | --- | --- |
| Structure | Range | *M* (SEM) | % of Total | *Z* | *p* |
| Frontal | 206 - 3155 | 1566 (238) | 45% | 3.180 | .001 |
| Parietal | 0 - 1230 | 402 (122) | 10% | 2.803 | .003 |
| Temporal | 0 - 2777 | 635 (216) | 15% | 2.803 | .003 |
| Occipital | 0 - 1692 | 644 (150) | 17% | 2.803 | .003 |
| Limbic | 0 - 839 | 80 (65) | 3% | 2.739 | .034 |

| B Gyrus Level |  |  |  |  |  |
| --- | --- | --- | --- | --- | --- |
| Structure | Range | *M* (SEM) | % of Total | *Z* | *p* |
| IFG | 0 - 1468 | 341 (152) | 11% | 2.023 | .022 |
| MeFG | 0 - 133 | 21 (12) | 1% | 2.023 | .022 |
| MFG | 0 - 1147 | 443 (114) | 13% | 2.803 | .003 |
| SFG | 0 - 962 | 404 (100) | 11% | 2.803 | .003 |
| PrG | 0 - 1034 | 294 (118) | 8% | 2.366 | .009 |
| PoG | 0 - 601 | 102 (48) | 2% | 2.201 | .014 |
| SPL | 0 - 682 | 94 (58) | 2% | 2.023 | .022 |
| ITG | 0 - 496 | 76 (46) | 2% | 2.201 | .014 |
| MTG | 0 - 2000 | 349 (163) | 8% | 2.201 | .014 |
| STG | 0 - 1367 | 231 (123) | 5% | 2.201 | .014 |
| IOG | 0 - 726 | 169 (67) | 5% | 2.023 | .022 |
| MOG | 0 - 1209 | 220 (99) | 6% | 2.521 | .006 |
| LgG | 0 - 371 | 83 (39) | 2% | 1.826 | .034 |
| Cun | 0 - 798 | 115 (65) | 3% | 1.826 | .034 |

Table. 1. Hits analysis of P3-like network at the Lobe (**A**), Gyrus (**B**), and Cell Type (**C**) levels reveal large overlap between the P3-like and P3 networks.
| C Cell Type Level |  |  |  |  |  |
| --- | --- | --- | --- | --- | --- |
| Structure | Range | *M* (SEM) | % of Total | *Z* | *p* |
| BA 10 | 0 - 1057 | 269 (118) | 7% | 1.826 | .034 |
| BA 9 | 0 - 525 | 198 (57) | 6% | 2.521 | .006 |
| BA 44 | 0 - 896 | 125 (72) | 4% | 1.826 | .034 |
| BA 8 | 0 - 1080 | 306 (120) | 11% | 2.023 | .022 |
| BA 6 | 0 - 1625 | 380 (169) | 11% | 2.201 | .014 |
| BA 4 | 0 - 344 | 87 (32) | 3% | 2.201 | .014 |
| BA 3 | 0 - 217 | 23 (17) | 1% | 1.826 | .034 |
| BA 7 | 0 - 1230 | 214 (107) | 6% | 2.201 | .014 |
| BA 40 | 0 - 909 | 118 (73) | 4% | 1.826 | .034 |
| BA 38 | 0 - 1205 | 206 (114) | 5% | 2.023 | .022 |
| BA 21 | 0 - 977 | 224 (104) | 6% | 2.023 | .022 |
| BA 20 | 0 - 589 | 113 (61) | 4% | 1.826 | .034 |
| BA 19 | 0 - 829 | 259 (89) | 7% | 2.366 | .009 |
| BA 18 | 0 - 774 | 282 (89) | 8% | 2.521 | .006 |
| BA 17 | 0 - 504 | 128 (49) | 4% | 2.023 | .022 |

Comparing the network underlying the P3-like components with those underlying P3 components reported in our companion paper [4], we found that the four lobes of the cerebral cortex are shared, indicating that both networks cover all lobes of the cerebral cortex. At the gyrus level, 12 out of the 14 gyri were shared **(Table. 1. shaded lines)**. P3-like components additionally involved the PoG and IOG while P3 components included the Pcun and FuG. At the cell type level, 11 of the 15 reported Brodmann areas were shared **(Table. 1. shaded lines)**. As expected, these regions cover all four lobes of the cerebral cortex.

Additional Brodmann areas unique to the P3-like network included BA44, BA4, BA3, and BA40 while the P3 network included BA46 and BA47. These differences may in part reflect the engagement of language areas of the brain under resting conditions and the engagement of more executive functions during task performance. Thus, the hits-based analysis revealed not only substantial overlap between the P3-like and P3 networks, but subtle differences between these highly similar networks, captured by measures of the network configuration.

Note that most of the average % of total hits values are fairly small (**Table. 1)**. This may reflect small contributions in all participants or large contributions by a few participants only, i.e. zero contributions by the rest of the participants. To disambiguate between these two possibilities, we show 3D plots of the % of totals for each individual and each lobe, with **Fig. 5A** highlighting the highly variable spatial configuration of the P3-like network across different individuals (different colors labeling different individuals) and **Fig. 5B** contrasting the more consistent cross-individual involvement of the frontal lobe with the highly variable involvement of temporal, parietal, and occipital lobes (different colors labeling different structures). These plots demonstrate that large individual differences can be revealed through this multi-dimensional hits-based analysis of P3-like components.

**Fig. 5.**
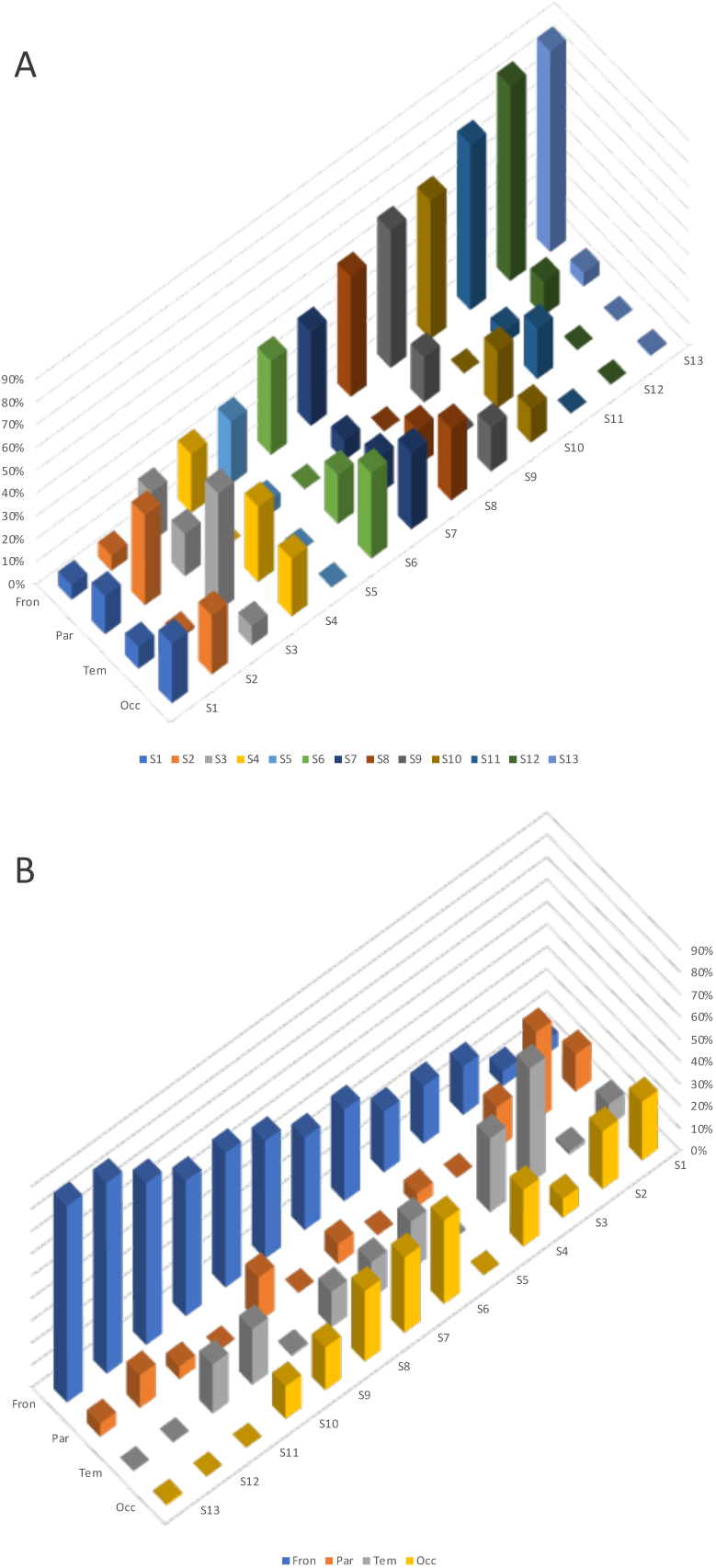
P3-like network configuration in all individuals. **(AB):** same data are plotted with different colors assigned to different individuals **(A)** and different structures **(B)**.

## IV. Conclusion

This present work shows that methodologically (1) a P3-like component can be reliably extracted by SOBI from each of the individuals’ (n=13) five minutes of resting-state EEG; and, (2) the spatial configuration of the neural network underlying this P3-like component can be localized using ECD models with high goodness of fit; (3) the network structure of the P3-like component can be quantitatively characterized for each individual through a hits-based analysis. Specifically, the spatial configuration of each individual’s P3-like network can be characterized as a vector of % of total hits whose length is defined by the number of constituent structures within the P3-like network and the values defined as the percentage of total hits that each of the constituent cortical structures make up for the whole network. We consider this P3-like network reflecting high temporal resolution synchrony of neuronal signaling across the cerebral cortex, which differs from the various networks reported by resting-state MRI work [32] reflecting synchronized BOLD signal changes, that are indirect measures of neuronal signaling. We note that although current work focuses on the characterization of the network spatial configuration, the more frequently used frequency domain analysis [7] can be used together to offer both spatial and temporal information in quantifying individual differences.

Because each SOBI component has a fixed scalp projection and a time course of activity, the P3-like component can be interpreted as reflecting synchronized neural network activity originating from multiple but spatially discontinuous structures. Through ECD source localization and hits-based analysis, we provided quantitative neuroanatomical characterization of the spatial configuration of the underlying neural generators. Similar to the neural network underlying the P3 component recovered from a visual oddball task, this P3-like network consists of cortical structures across all four lobes of cerebral cortex, matching well with the broad innervation of the neocortex by the major neuromodulatory systems. Therefore, we consider the quantitative characteristics of the P3-like network recoverable from resting-state brain as neural computationally relevant parameters, thus potentially good candidates for being effective neuromarkers.

Furthermore, because the traditional use of P3 amplitude and latency are sensitive to stimulus intensity, frequency, inter-trial interval, and past history of experiencing similar stimuli, neuromarkers based on characteristics derived from task-free and resting-stating EEG are, in principle, more likely to capture trait-like individual differences, not “contaminated” by variations associated with these other factors. Similar to the study of the resting-state brain activity using fMRI, here, the quantitative characterization of the P3-like network from resting-state EEG data also has all the benefits of not requiring tasks to be performed by the individual under investigation. Different from the resting-state MRI studies, the resting-state EEG studies are substantially less expensive and more convenient for both the study participants and the investigators. This increased feasibility of obtaining the P3-like network parameters as neuromarkers from resting-state EEG also affords a wider range of applications beyond biomedical research and clinical treatment, to include basic research in understanding the brain in natural context.

The hits-based analysis method, recently introduced by our group (see companion paper [4]), offered a novel approach to quantitatively characterize the spatial configuration of a distributed network at the level of individuals. With this method, each individual can be described by a vector, whose length is determined by the number of all observed grey matter brain structures across all participants, and whose elements are defined by each of the structure’s hits contribution to the total grey matter hits in that individual. In such a high dimensional space, reliability or variability of each brain structure or each subnetwork in this P3-like global network can be observed and quantified. In such a space, individual differences in any of its sub-spaces can potentially reveal clusters associated with different disease conditions or different cognitive and emotional regulation capacities. The present work lays the methodological foundation for future studies of individual differences in health and disease and in both experimental and natural contexts.

## Acknowledgment

This work was supported by a grant from the University of Hong Kong [104004683] and a donation from the Sweeting Memorial Fund awarded to A.C.T. We thank Drs. R. Sun and E. Tsang and X. Niu for their assistance.

